# Mechanical cues trigger phellem differentiation during barrier transition

**DOI:** 10.64898/2026.03.10.710754

**Authors:** Jennifer López-Ortiz, Damien De Bellis, Etienne Bellani, Niko Geldner, Juan Alonso-Serra, Hiroyuki Iida, Ari Pekka Mähönen

**Affiliations:** Department of Organismal and Evolutionary Biology, Faculty of Biological and Environmental Sciences and Viikki Plant Science Centre, University of Helsinki, 00014 Helsinki, Finland; Department of Plant Molecular Biology, University of Lausanne, Lausanne, Switzerland; Electron Microscopy Facility, University of Lausanne, Lausanne, Switzerland; Umeå Plant Science Center, Department of Forest Genetics and Plant Physiology, Swedish University of Agricultural Sciences, 90736, Umeå, Sweden

## Abstract

A continuous and uninterrupted barrier tissue is essential for protecting internal tissues from the external environment. In many dicot roots, the endodermis fulfils this role during primary growth; however, the onset of secondary growth in the vascular tissue coincides with the rupture of the overlying endodermis and formation of a new barrier, the periderm, beneath it. Despite the importance of barrier integrity for plant survival, the mechanisms coordinating this barrier transition have remained unexplored. Here, we show that endodermal rupture in *Arabidopsis thaliana* roots releases mechanical constraints on the underlying properiderm. This release leads to cell expansion and triggers differentiation of the outermost properiderm cells into phellem, a ligno-suberized barrier cell type. Premature relief of mechanical constraints, through hyperosmotic stress or targeted endodermal ablation, is sufficient to trigger phellem differentiation. FERONIA, a cell wall integrity receptor kinase, is required for phellem differentiation in response to mechanical stress. Overall, our findings establish mechanical stress as an instructive cue for phellem differentiation, revealing how plants convert mechanical signals into robust, adaptive protective barriers.

---

During primary root growth, the endodermis forms the critical apoplastic barrier that protects the stele from biotic and abiotic stresses. This barrier is established through the lignification of the Casparian strip and by suberin deposition in the endodermal cell walls^1–3^. In most seed plants, this barrier tissue is disrupted as roots undergo secondary growth, when radial expansion of the stele generates additional vascular tissues to enhance transport efficiency and structural support^4^. In *Arabidopsis* roots, secondary growth initiates with procambial divisions followed by pericycle divisions. The vascular cambium and phellogen (cork cambium) are established from procambium and pericycle cells^5–7^, respectively, with Xylem Pole-associated Pericycle (XPP) cells contributing to both meristems^7^. As the expanding stele ruptures the overlying tissues, the pericycle lineage forms the functional periderm –a protective tissue composed of the meristematic phellogen, inward-produced phelloderm, and outward-produced differentiated phellem (Fig. 1a). The phellem differentiates by depositing ligno-suberized secondary cell walls and replaces the endodermis as the outermost barrier to restrict water and gas diffusion and to prevent pathogen entry^8–10^. Here, we define the developmental window encompassing the switch from endodermis to phellem as the *barrier transition*. This process must be tightly coordinated to maintain continuous protection of the stele, yet the mechanisms controlling its timing and identity shifts remain largely unknown.

**Fig. 1.**
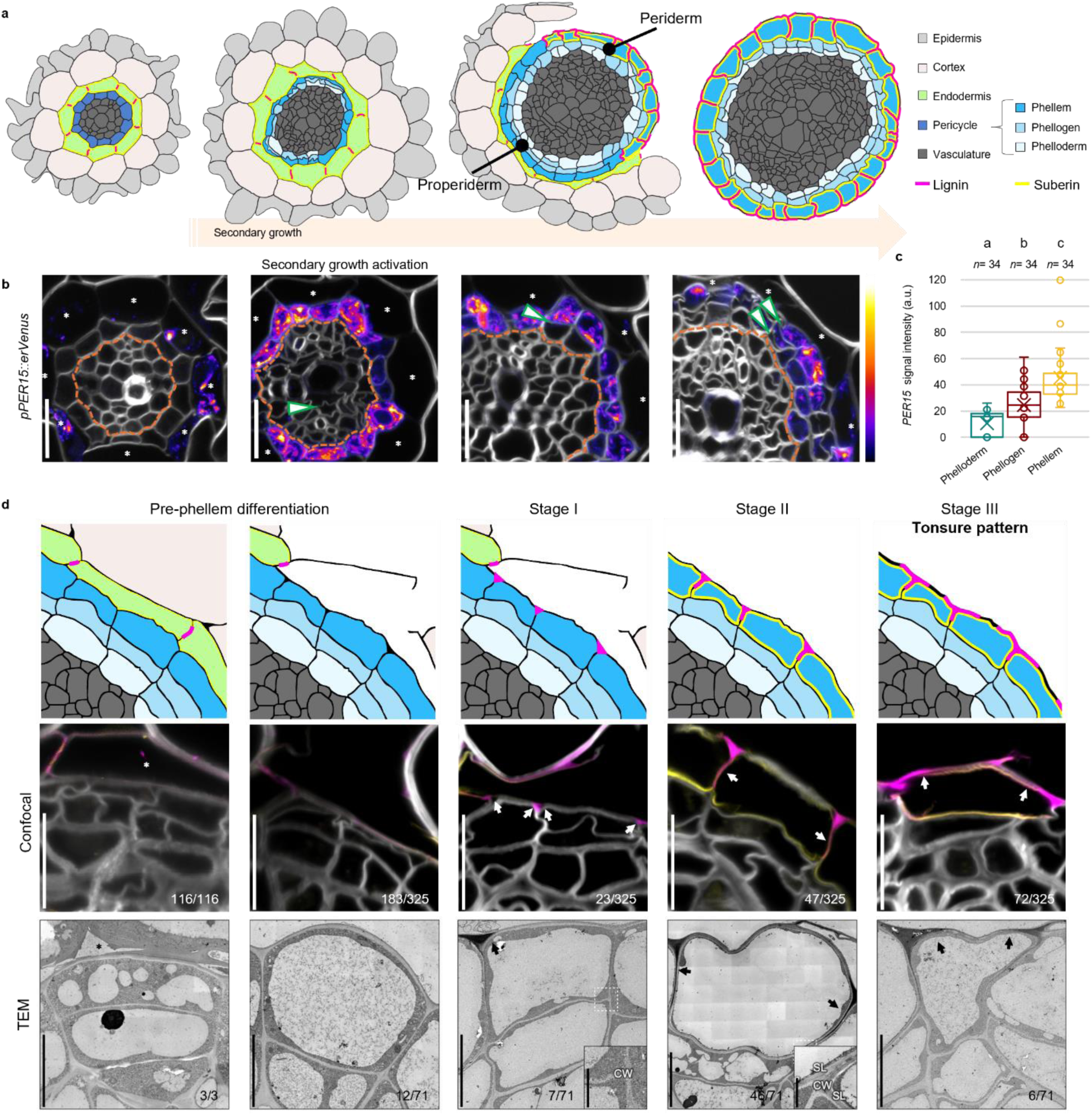
Phellem identity is specified upon secondary growth activation and differentiation occurs after endodermal rupture. **a**, Schematic root cross-sections illustrating the transition from primary to secondary growth. **b**, Confocal microscopy of the *pPER15::erVenus* reporter line across developmental stages. Venus signal intensities are shown according to the colour map on the right, from high (top) to low (bottom). Green arrowheads indicate periclinal divisions. **c**, Quantification of *PER15* signal intensity in three-cell radial files. Boxes represent the median and interquartile range (IQR); whiskers extend to 1.5×IQR; x, the mean; circles, individual cell measurements. One-way ANOVA followed by Bonferroni-corrected two-tailed Wilcoxon rank-sum test were used; different letters indicate significant differences between groups (*P* < 0.001). n indicates the number of cells analyzed. **d**, Wild-type root cross-sections showing the phellem differentiation stages based on lignin and suberin deposition. Top, schematic illustrations; middle, confocal microscopy; bottom, TEM. TEM close-up images of walls of exposed cells at Stage I and II showing the structure of suberin lamellae (SL) and the primary cell wall (CW). Fractions on the panel indicate the proportion of cells displaying similar lignin and suberin patterns when endodermis is intact (leftmost column) or ruptured (other columns). White and black arrows indicate lignified regions in confocal and TEM images, respectively. Endodermal cells are marked with asterisks in **b** and **d**. Cell walls were stained with SR2200 (grays); lignin with Basic Fuchsin (magenta); and suberin with Fluorol Yellow (yellow). Experiments in **b–c** were independently repeated three times; confocal panels in **d**, five times; and the TEM panels **d**, four roots from two independent experiments. Scale bars: 20 µm (**b**), 10 µm (**d**, middle), 5 µm (**d**, bottom), 1 µm (d, bottom; close-ups).

In this study we show that pericycle cells acquire periderm identity prior to their first divisions during secondary growth, yet their differentiation into phellem is tightly correlated with the rupture of the overlying endodermis. Neither direct environmental exposure nor endodermal identity and barrier function are required for phellem differentiation. Instead, changes in mechanical patterns are sufficient to trigger premature periderm identity and phellem differentiation in the pericycle lineage, revealing that the barrier transition from endodermis to phellem is orchestrated primarily through mechanotransduction. Mechanical stress is sensed through a competent cell wall, inducing cellular deformation, and is transduced through the receptor kinase FERONIA (FER) to trigger phellem differentiation.

## RESULTS

### Phellem differentiation correlates with endodermis rupture

To characterise phellem establishment, we analysed cell division patterns in the pericycle of wild-type roots. Following the first periclinal division in the pericycle cell, we found that the outer daughter cell frequently further divided periclinally, resulting in a radial cell file consisting of three cells (269 out of 321 pericycle lineages, Extended Data Fig. 1a). These results indicate that the first pericycle division establishes the phellogen, which initially produces the phelloderm, while a subsequent division within the phellogen gives rise to undifferentiated phellem (Extended Data Fig. 1a). Because phellem has not yet differentiated at this stage, and the periderm therefore does not yet function as a barrier, we refer to this early pre-differentiation stage as the *properiderm* (Fig. 1a).

To determine when the phellem identity is acquired in the properiderm, we analysed the phellem-expressed reporter lines in which endoplasmic-reticulum localised Venus (erVenus) is expressed under the control of *PEROXIDASE 15* (*PER15*) or *PER49* promoter^9,11,12^. These reporter lines showed fluorescence signals in the endodermis prior to secondary growth initiation (Fig. 1b; Extended Data Fig. 1c). When secondary growth is activated, we detected that the expression of these reporter lines shifted from the endodermis to the pericycle, and their expression peaked in the outermost cells after the cells in the pericycle lineage (i.e. in the properiderm) underwent periclinal divisions (Fig. 1b-c; Extended Data Fig. 1b-d). These results indicate that phellem identity is gradually specified before pericycle cells divide and phellem/phellogen identity is subsequently confined to the outer domain of the properiderm.

Next, we examined phellem differentiation based on lignin and suberin deposition. The formation of secondary cell wall was visualised using Basic Fuchsin and Fluorol Yellow to stain lignin and suberin, respectively. Secondary cell wall in the phellem was not detected while the overlying endodermis remained intact (116 out of 116 cells, Fig.1d). Lignin and suberin staining were found in the outermost properiderm cells after the cell wall of the overlying endodermis was ruptured (142 out of 325 cells, Fig.1d). Endodermis rupture occurred predominantly in the Phloem Pole-associated Pericycle (PPP) region (35 out of 58 roots, Extended Data Fig. 1e-f). Less frequently in the XPP or Non-Pole-associated Pericycle (NPP) regions (12 and 11 out of 58 roots, respectively, Extended Data Fig. 1e-f). Phellem occasionally differentiated from the outer cell of a two-cell file properiderm when the overlying endodermis was prematurely disrupted (37 out of 105 lineages, Extended Data Fig. 1g). Thus, we speculated that the disruption of the overlying endodermis may provide cues to activate phellem differentiation. Detailed analysis revealed that lignin staining first appeared at outer cell-corners (Stage I, 23 out of 325 cells) and then extended along radial walls coinciding with uniform suberin deposition (Stage II, 47 out of 325 cells). Finally, lignin was detected in the outer tangential wall while the central domain remained unlignified, forming a distinctive *tonsure*-like pattern (Stage III, 72 out of 325 cells, Fig. 1d). Transmission Electron Microscopy (TEM) confirmed these stages, revealing electron-dense lignin depositions and lamellar suberin accumulation from Stage II onwards (Fig. 1d). The tonsure pattern persisted in the mature phellem of 20-day-old roots, indicating that it is a stable characteristic of mature phellem (Extended Data Fig. 1h).

### Pericycle is competent to differentiate into phellem

Given the correlation between the deposition of the secondary cell wall polymers and the disruption of the overlying endodermis, we tested whether the premature endodermis removal induces phellem differentiation in the pericycle. To this end, we surgically ablated the outer tissues with a needle and exposed pericycle cells in the mature root region of 4-day-old seedlings, before the activation of secondary growth (Fig. 2a). At 3 hours after injury (hai), we detected prematurely *pPER15::erVenus* signals in the pericycle cells (Fig. 2b-c). This shift normally occurs later in development, after the activation of secondary growth (Fig. 1b). By 48 hai, exposed pericycle cells frequently divided periclinally (55 out of 84 cells) and the outer cells deposited lignin and suberin following the stages of native phellem differentiation (Fig. 2d-f). In contrast, control pericycle cells did not undergo cell division and showed no lignin or suberin deposition (Fig. 2g; mock). TEM analysis confirmed that, by 72 h, lignin and suberin deposition in these cells corresponded to Stage III of the native phellem development (Fig. 2e). These results demonstrate that ablation of outer tissues induces premature differentiation of pericycle into phellem.

**Fig. 2.**
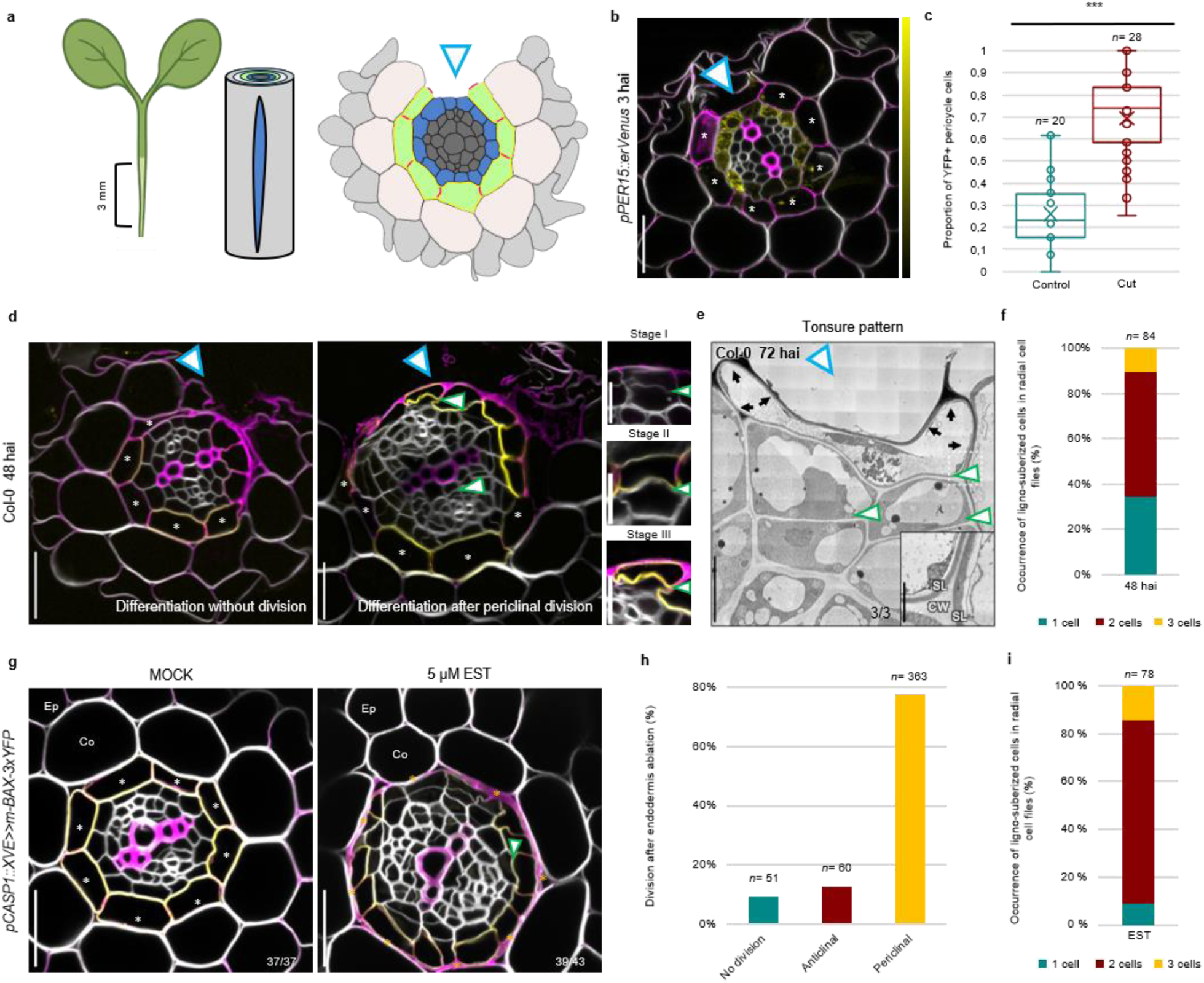
Endodermis ablation triggers premature phellem differentiation in pericycle cells. **a**, Schematic of the surgical injury experiment. Four-day-old roots were longitudinally wounded within 3 mm below the root-hypocotyl junction to remove pericycle-overlying tissues. **b**, Confocal microscopy of *PER15::erVenus* roots at 3 hai. Venus signal intensities are shown according to the colour scale on the right. **c**, Quantification of the proportion of YFP-positive cells in control (Control) and injured roots (Cut). Boxes represent the median values and interquartile range (IQR), whiskers extend to 1.5×IQR, x marks the mean, and the circles indicate measurements from individual roots. Two-tailed Welch’s t-test was used (***, *P* < 0,001). n indicates the number of examined roots. **d–e**, Confocal microscopy (**d**) or TEM image (**e**) of lignin and suberin deposition patterns in wild-type root at 48 and 72 hai. TEM close-up showing the structure of suberin lamellae (SL) and the cell wall (CW). **f**, Quantification of ligno-suberized cells in radial cell files at 48 hai, indicating the frequency of phellem differentiation. n indicates the number of examined lineages. **g**, Confocal microscopy of lignin and suberin deposition patterns in *pCASP1::XVE>>m-BAX-3xYFP* roots grown on mock (MOCK) or 5µM EST (5µM EST) plate for 6 days. Fractions on the panel indicate the proportion of roots exhibiting the phenotypes as shown in the panel. Collapsed endodermal cells are indicated with orange asterisks. Ep, epidermis; Co, cortex. **h**, Quantification of pericycle division orientation after endodermal collapse (EST treatment). n indicates the number of examined divisions. **i**, Quantification of ligno-suberized cells in radial cell files in EST treated roots, indicating the frequency of phellem differentiation. n indicating the number of examined lineages. Endodermal cells are marked with white asterisks in **b**,**d**,**g**. Green arrowheads, periclinal cell division (**d**,**e**,**g**); blue arrowheads, wound sites (**a,b,d,e**). Cell walls were stained with SR2200 (grays); lignin with Basic Fuchsin (magenta); and suberin with Fluorol Yellow (yellow). Experiments in **b–c** were repeated twice (independently from control, four times); **d**,**f,** three times; **e,** three cells from one root; **g–i,** four times. Scale bars: 20 µm (**b**,**d**,**g**), 5 µm (**e**), 1 µm (**e**; close-up).

To test whether direct exposure of pericycle to the outside environment is required for phellem differentiation, we created a genetic ablation system to specifically target the endodermis. We conditionally triggered programmed cell death in endodermal cells by inducing the expression of *murine-BAX-3xYFP*^13^ (*mBAX-3xYFP*) under the *CASPARIAN STRIP MEMBRANE DOMAIN PROTEINS1* (*CASP1*) promoter upon estradiol treatment (*pCASP1::XVE>>mBAX-3xYFP*). The *CASP1* promoter is active in the differentiating endodermis where the Casparian strip forms^14^. We treated *pCASP1::XVE>>mBAX-3xYFP* seedlings with estradiol upon germination and analysed the mature part of their roots six days later. We observed that the ablated endodermal cells collapsed, showing internal Basic Fuchsin staining, and the underneath pericycle cells divided periclinally (Fig. 2 g-i), consistent with previous reports showing that laser ablation of endodermis induces cell division in underlying pericycle^15,16^. Furthermore, we found that these outermost properiderm layer showed the lignin and suberin patterns of native phellem (Fig. 2g). Altogether, phellem differentiation occurred even when the overlying epidermis and cortex remained intact, suggesting that direct environmental exposure is not required for phellem differentiation.

### Endodermis identity is not required to repress phellem differentiation

Since direct environmental exposure was not required for phellem differentiation, we hypothesised it is instead triggered by the removal of either (1) a genetic inhibitory signal or (2) a mechanical constraint from the endodermis.

Endodermis identity can influence neighbouring cell fates through intercellular signals, as in vascular patterning^17,18^. To test the role of endodermis-derived genetic factor inhibiting phellem differentiation, we analysed *shortroot-2* (*shr-2*) mutant roots, which forms a single ground-tissue layer lacking endodermis identity^19–21^. If such a signal exists, *shr-*2 pericycle cells should differentiate precociously into phellem, but no lignin or suberin was detected prematurely in pericycle cells before or after secondary growth activation (26 out of 26 and 57 out of 60 roots, respectively) (Fig. 3a). Although *shr-2* roots showed severe growth defects, cells in pericycle lineage are competent to differentiate into phellem, as mature 16-day-old *shr-2* roots displayed differentiated phellem and surgical ablation of 4-day-old *shr-2* roots triggered lignin and suberin deposition in exposed properiderm within 48 hai (27 out of 30 roots) (Extended Data Fig. 2a; Fig. 3b). We also analysed the other endodermis-related mutants with impaired barrier function, including *scarecrow-4* (*scr-4*)^21,22^, *myb36-2;schengen3-3* (*myb36-2;sgn3-3*) double mutant^23^, and the quintuple mutant of *GDSL-type esterase/lipase proteins 22, 38, 49, 51, 96* (*gelpQ*)^24^. Similar to *shr-2* mutant and Col-0, none of these mutants showed premature differentiation of the cells in the pericycle lineage (Extended Data Fig. 2b). Altogether, these results indicate that neither endodermal identity nor its barrier function are required to inhibit phellem differentiation.

**Fig. 3.**
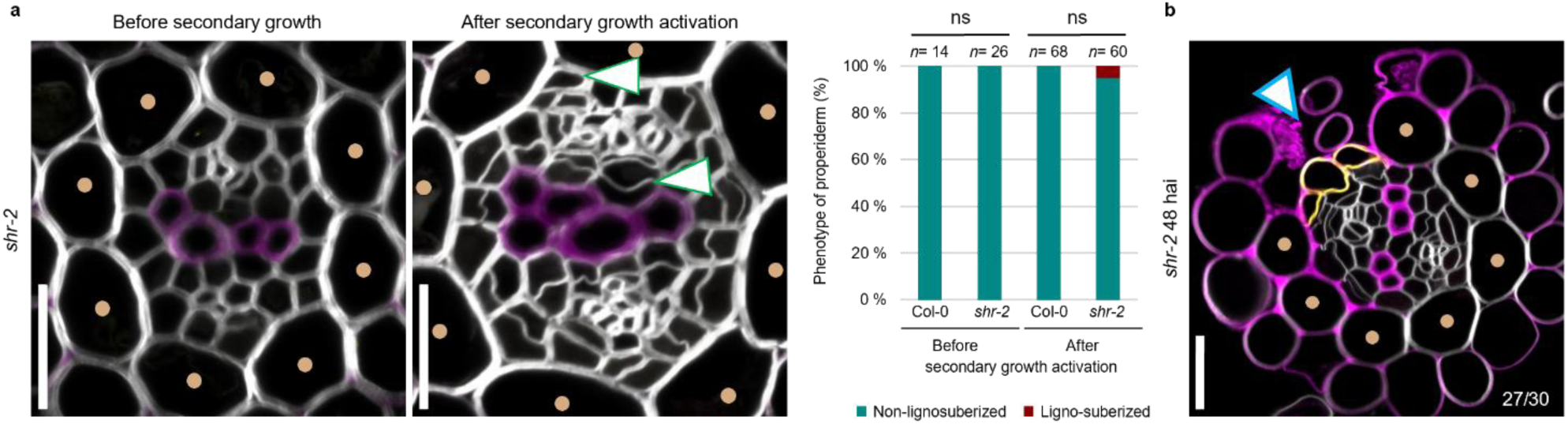
Lack of endodermal identity does not trigger premature phellem differentiation. **a**, Confocal microscopy of endodermis-defective *shr-2* mutant roots before and after secondary growth activation, showing the lignin and suberin deposition patterns (left panels), and quantification of the proportion of roots showing phellem differentiation in the pericycle cells (right plot). Two-sided Fisher’s exact test was used to compare Col-0 and *shr-2* phenotypes before and after secondary growth activation independently (n.s., *P ≥* 0.05). Premature pericycle cell divisions were observed in *shr-2* roots. Green arrowheads indicate periclinal cell divisions. **b**, Confocal cross-section of a *shr-2* root at 48 hai showing lignin and suberin deposition patterns. Fraction indicates the proportion of roots showing secondary cell wall deposition in exposed pericycle cells. Blue arrowhead, the injury site; light brown dots, the mutant ground tissue layer in *shr-2* mutant. Cell walls were stained with SR2200 (grays); lignin with Basic Fuchsin (magenta); and suberin with Fluorol Yellow (yellow). Experiments in **a** were repeated four times and in **b** twice. Scale bars: 20 µm (**a**,**b**).

### Release of mechanical constraints drives cellular expansion and triggers phellem differentiation

Next, we tested the second hypothesis: mechanical cues regulate phellem differentiation. We observed a local expansion of the properiderm, specifically at sites where the adjacent endodermis was ruptured. Quantitative analysis of cell size during normal development revealed that outermost properiderm cells or the whole radial cell file (i.e. lineage) were larger when exposed compared to non-exposed cells, implying that the overlying tissues impose a mechanical constraint, and their rupture leads to changes in the mechanical environment causing cell deformation (Fig. 4a-b; Extended Data Fig. 3a). Notably, cell expansion among exposed outermost properiderm cells was heterogeneous, with ligno-suberized cells exhibiting significantly greater enlargement than exposed cells that did not differentiate (Fig. 4b). To investigate the temporal relationship between properiderm cell deformation and the formation of secondary cell wall, we surgically ablated the outer tissues of the mature region of 4-day-old roots to expose pericycle cells. At 3 hai, the exposed PPP cells and NPP cells were already significantly larger than their non-exposed counterparts, and their size increased over time (Fig. 4c-d). At 24 hai, the first divisions were evident, and the exposed cell occasionally differentiated into phellem, predominantly at Stage I, where the exposed region showed continuous or patchy phellem differentiation (8 and 11 out of 30 roots, respectively) (Extended Data Fig. 3b-c; Fig. 4e). By 48 hai, the outer cells formed a continuous ligno-suberized Stage III phellem layer (25 out of 29 roots) (Fig. 2d and 4e). Taken together, the rupture of overlying tissues rapidly caused the expansion of the underlying pericycle cells followed by periclinal divisions and the deposition of secondary cell wall, suggesting that the release of mechanical constraint triggers cell deformation and the phellem differentiation.

**Fig. 4.**
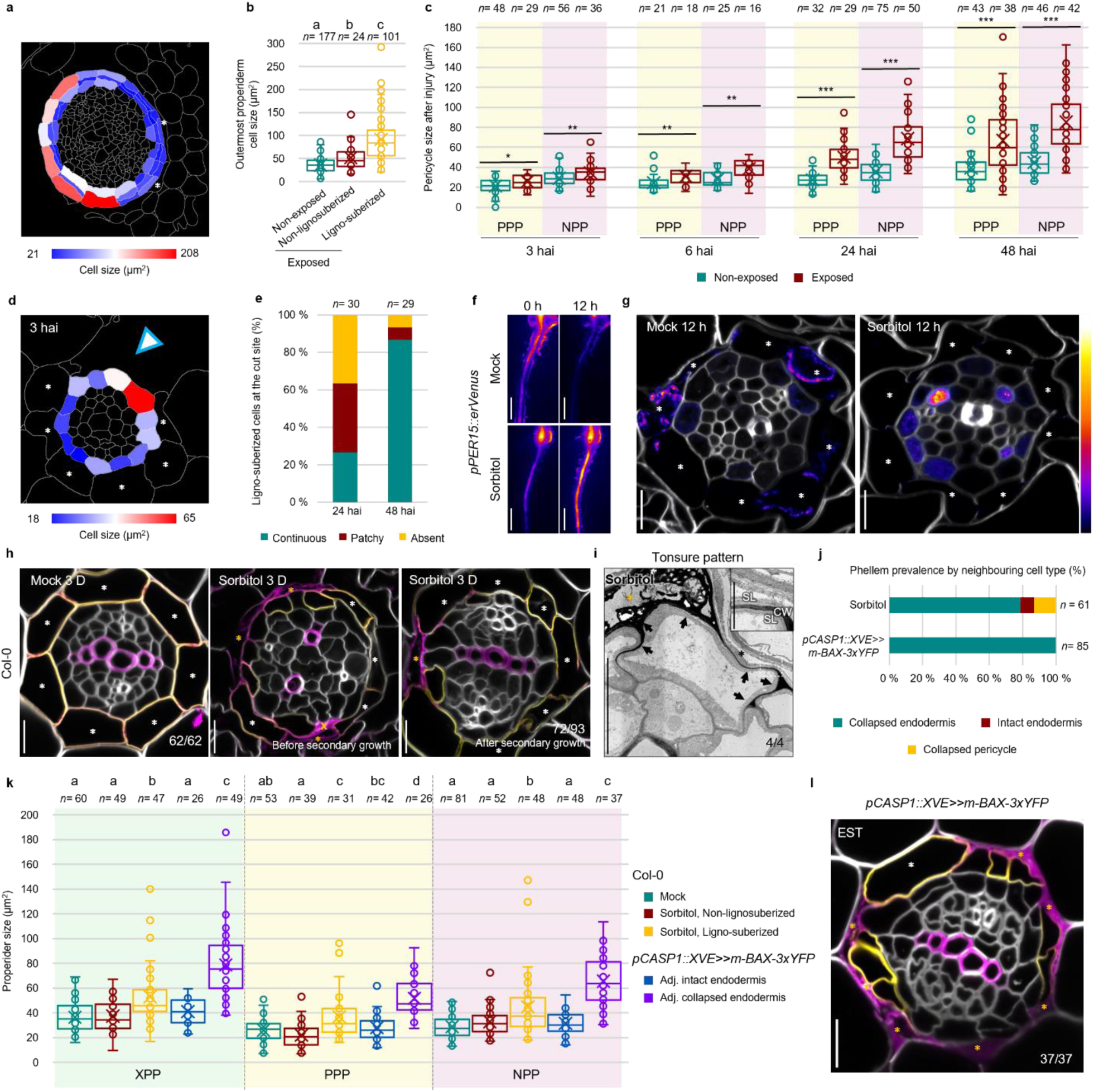
Release of endodermal mechanical constraint promotes pericycle expansion and phellem differentiation. **a**, Heatmap of properiderm cell area in wild-type roots at the barrier transition stage. **b**, Quantification of outermost pericycle-derived cell area at three distinct stages: the outermost cell overlain by endodermal cells (Non-exposed), exposed to the surface without (Non-lignosuberized) or with (Ligno-suberized) lignin and suberin deposition. **c**, Quantification of PPP and NPP cell or lineage area over time after surgical injury. **d**, Heatmap of pericycle cell size at 3 hai. Blue arrowhead marks the cut site. **e**, Ligno-suberized cell coverage at the cut site at 24 and 48 hai. **f–g**, *pPER15::erVenus* seedlings treated with mock or sorbitol for 12 h showing *PER15* signal under stereomicroscopy (**f**) and confocal microscopy (**g**). Venus signal intensities are shown according to the colour map on the right. **h–i**, Wild type roots 3 days after mock or sorbitol treatment showing lignin and suberin patterning by confocal microscopy (**h**) and TEM (**i**). TEM close-up show suberin lamellae (SL) and primary cell wall (CW). Orange crosses indicate collapsed pericycle cells. **j**, Phellem prevalence relative to neighbouring cell type state in sorbitol-treated wild-type roots and EST-treated *pCASP1::XVE>>m-BAX-3xYFP* roots. **k**, Pericycle cell area in mock- and sorbitol-treated wild-type roots and EST-treated *pCASP1::XVE>>m-BAX-3xYFP* roots. **l**, Confocal microscopy of EST-treated *pCASP1::XVE>>m-BAX-3xYFP* roots showing partial endodermis ablation. Fractions in panels denotes the frequency of the presented phenotype in different roots (**h**,**l**) or cells (**i**). Boxes represent median and interquartile range (IQR), whiskers extend to 1.5×IQR, x marks the mean, and the circles indicate individual measurements. In **b** and **k**, one-way ANOVA followed by Bonferroni-corrected two-tailed Wilcoxon rank-sum test were used; different letters indicate significant differences between groups (*P* < 0.05). In **c**, two-tailed Welch’s t-test was used to compare PPP and NPP cell size at 3 hai and PPP at 24 hai; and two-tailed Wilcoxon rank-sum test was used to compare the remaining (*, *P* < 0,05; **, *P* < 0,01; ***, *P* < 0,001). n indicates the examined cells (**b,j,k,** hai in **c**) or lineages (24 and 48 hai in **c**) or roots (**e**). Intact endodermal cells are marked with white asterisks in **a**,**d**,**g**,**h**,**l** and in black asterisks in **i**. Collapsed endodermal cells are indicated with orange asterisks in **h**,**i**,**l**. Cell walls were stained with SR2200 (white); lignin with Basic Fuchsin (magenta); and suberin with Fluorol Yellow (yellow). Experiments in **a–g** were independently repeated three times; Col-0 samples in **h**,**j**,**k** were repeated three times; **i** four cells were examined from a single root. Experiments using *pCASP1::XVE>>m-BAX-3xYFP* in **j-l** were repeated four times. Scale bars: 1 mm (**f**), 10 µm (**g**,**h**,**l**), 5 µm (**i**), 500 nm (**i**; close-up).

To test whether changes in mechanical constraint are sufficient to induce phellem identity, we treated 3-day-old *pPER15::erVenus* and *pPER49::erVenus* seedlings with 400 mM sorbitol. This hyperosmotic treatment leads to cell water efflux, turgor pressure reduction, and ultimately cell deformation^25,26^. Within 12 hours, the *PER15* and *PER49* expressions were upregulated, and their expression pattern in the endodermis shifted predominantly to the pericycle (Fig. 4f-g; Extended Data Fig. 4a), suggesting premature phellem identity acquisition. Notably, at this early time point, pericycle cell size was not yet different between mock and sorbitol-treated roots, suggesting that the phellem identity activation occurs prior to detectable cell deformation (Extended Data Fig. 4b). Three days after the sorbitol treatment, we found that some pericycle cells exhibited distinct stages of secondary cell wall deposition similar to those of native phellem (72 out of 93 roots; Fig. 4h). TEM analysis also confirmed the tonsure pattern and suberin lamellae deposition of these pericycle cells (Fig. 4i; Extended Fig Data 4c), indicating that hyperosmotic treatment induces phellem-like features in pericycle cells. While hyperosmotic stress can activate the abscisic acid (ABA) signalling pathway^27^, which is known to regulate suberization in the endodermis^28^, we found 1 µM ABA treatment did not increase *PER15* and *PER49* expression (12 h post-treatment) or induce the formation of the secondary cell wall in the pericycle, even when we increased the concentration to 5 µM of ABA (3 days post-treatment) (Extended Data Fig. 4a,d-g). Moreover, lignin and suberin staining was still detected in the pericycle of the ABA insensitive mutants, *aba insensitive1-1* (*abi1-1*)^29^ and *pyrabactin resistance1*;*pyr1-like1,2,4,5,8* (*pyr1;pyl1,2,4,5,8*)^30^, after the sorbitol treatment (Extended Data Fig. 4i). Although applied ABA reduces turgor pressure^31^, only sorbitol triggers phellem differentiation. While analysing this discrepancy, we noticed that sorbitol-treated roots exhibited extensive endodermal cell collapse (199 collapsed out of approximately 750 endodermal cells), detected by internal Basic Fuchsin staining and confirmed by TEM (Fig. 4h-i; Extended Data Fig. 4c), reflecting the heterogeneous susceptibility of cells to hyperosmotic stress^32^. Unexpectedly, stele enlargement was observed only in sorbitol-treated roots, not in mock or ABA treatment (Extended Data Fig. 4k). Under hyperosmotic conditions, endodermal cell collapse may relieve local mechanical constraints, allowing underlying cells to expand and differentiate into phellem, consistent with reports that turgor loss in dead cells drives neighbouring cells expansion toward the wound^33,34^. Accordingly, fully differentiated phellem cells were predominantly adjacent to collapsed endodermal cells (48 out of 61 cells), whereas differentiation was rare underneath living endodermal cells (5 out of 61 cells) or besides collapsed pericycle cells (8 out of 61 cells) (Fig. 4j). Moreover, differentiated phellem cells were significantly larger than non-differentiated cells across pericycle lineages, further supporting the idea that cellular expansion triggers phellem differentiation (Fig. 4k).

Given that hyperosmotic stress compromises multiple cell types and pericycle is particularly susceptible (132 collapsed out of approximately 1100 pericycle cells), we applied targeted, genetic ablation of endodermis (*pCASP1::XVE>>mBAX-3xYFP*) to specifically address the role of endodermis in the mechanical stress. Ablation and subsequent collapse of endodermal cells resulted in stele enlargement (Extended Data Fig. 4k). In partially ablated regions, pericycle cells beneath collapsed endodermal cells were significantly larger than those beneath alive cells (Fig. 4k-l). Strikingly, phellem differentiation occurred only beneath ablated endodermal cells (Fig. 4j). Altogether, these results demonstrate that the endodermis functions as a restrictive mechanical barrier, and its removal relieves physical constraint in inner tissues, thereby enabling phellem differentiation.

### FER monitors cell wall integrity to trigger phellem differentiation under mechanical stress

Release of mechanical constraints following endodermis rupture likely generates local stress on properiderm cells that might be experienced by the cell wall since it is the principal load-bearing structure^35^. Cellulose provides tensile strength and pectin remodelling modulates stiffness and extensibility^36^. To test whether cell wall mechanics are required for phellem differentiation, we disrupted cell wall integrity (CWI) during the barrier transition stage. We altered cell wall mechanics by inhibiting cellulose synthesis with Isoxaben (ISX)^37,38^ or pectin methylesterase activity with Epigallocatechin gallate (EGCG) to increase pectin methyl-esterification^39,40^. After one day of treatment, overall *PER15* and *PER49* expression decreased, mildly with ISX and strongly with EGCG, with *PER15* showing a pronounced decrease in the directly exposed outermost cell (Fig. 5a-d; Extended Data Fig. 5a-b). This reduction correlated with fewer ligno-suberized cells upon these treatments, shifting from continuous to patchy or absent coverage (Fig. 5e-g). These results suggest to a potential role for CWI in initiating and sustaining phellem differentiation.

**Fig. 5.**
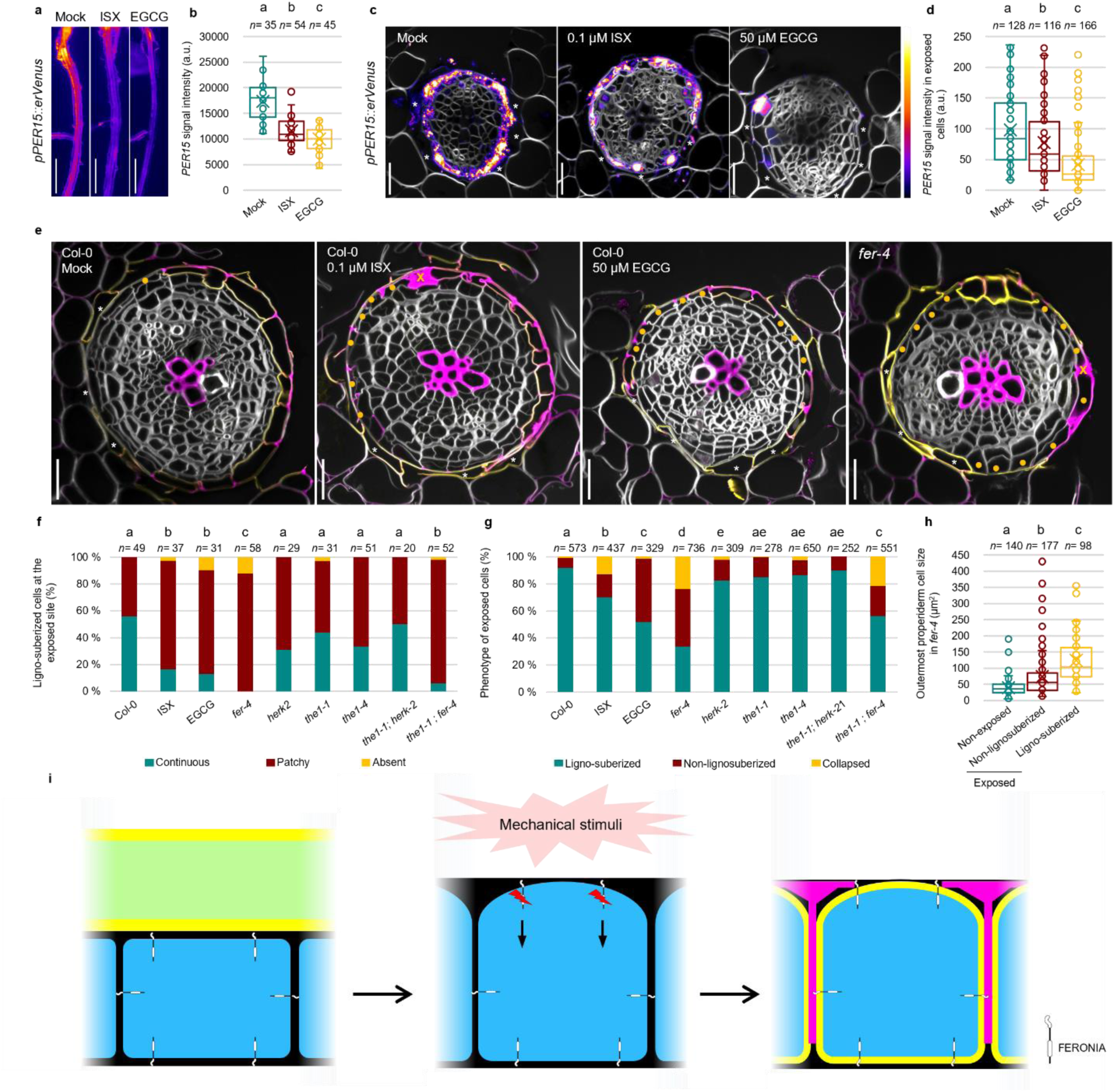
**FER-mediated mechanosensing drives phellem differentiation in response to mechanical stress. a–d**, *pPER15::erVenus* signals in their roots treated with mock (Mock), isoxaben (ISX) or EGCG (EGCG) for 24 h during the transition stage (**a**) and the quantification of the overall *PER15* signal intensity (**b**). Confocal microscopy of the *pPER15::erVenus* treated with mock (Mock), isoxaben (ISX) or EGCG (EGCG) for 24 h during the transition stage (**c**) and quantification of the signal intensity in exposed properiderm cells (**d**). Venus signal intensities are shown according to the colour map on the right. **e–g**, Wild-type seedlings treated with mock, ISX or EGCG for 24 h and 11-day-old CWI mutants at the barrier transition stage, stained for lignin and suberin. **e**, Confocal microscopy of representative phenotypes in wild-type roots treated with mock, isoxaben, and EGCG and *fer-4* mutant roots, stained for lignin and suberin. Orange dots mark undifferentiated phellem exposed cells; orange crosses indicate collapsed cells. Quantification of the barrier continuity (**f**) and cell-phenotype coverage (**g**) at the exposed region. **h**, Quantification of outermost properiderm cell area in *fer-4* when surrounded by endodermal cells or exposed, with or without phellem differentiation. **i**, Schematic of FER-dependent mechanosignalling triggering phellem differentiation after endodermis rupture. In **b**,**d,** and **h**, boxes represent median and interquartile range (IQR), whiskers extend to 1.5×IQR, x marks the mean, and the circles indicate individual measurements. In **b**, one-way ANOVA followed by Bonferroni-corrected pairwise Welch’s t-tests was used; in **d** and **h**, one-way ANOVA test followed by Bonferroni-corrected Wilcoxon rank-sum test was used; and **f–g** pairwise Fisher’s exact tests with Bonferroni correction were used to compare phellem counts among genotypes (different letters indicate significant differences between groups (*P* < 0.05). Endodermal cells are marked in asterisks. Cell walls were stained with SR2200 (white); lignin with Basic Fuchsin (magenta); and suberin with Fluorol Yellow (yellow). Phenotypes of *fer-2*, *herk2*, *the1-1* and *the1-1;herk2* were independently examined six times, whereas those of *the1-4* and *the1-1;fer-4* were examined four times. Scale bars: 1 mm (**a**) 20 µm (**c**,**e**).

To explore whether specific CWI sensors control phellem differentiation, we analysed mutants of the CrRLK1L receptor-like kinases family, including FERONIA (FER), THESEUS1 (THE1), HERKULES2 (HERK2). While *herk2*, *the1-1*, *the1-4* and *the1-1;herk2*^41–43^ resembled wild type phenotypes, *fer-4* mutant^44^ failed to form a continuous ligno-suberized layer (Fig. 5e-g; Extended Data Fig. 5a). Additionally, the outermost properiderm cells in *fer-4* occasionally collapsed (Fig. 5e-g; Extended Data Fig. 5c). Quantitative analysis of cell size in *fer-4* showed that cell expansion occurs normally upon exposure, yet differentiation is impaired, indicating that FER-mediated mechanotransduction is essential for ligno-suberin deposition (Fig. 5h). The *the1-1;fer-4* double mutant^45^ showed an intermediate phenotype between wild-type and *fer-4* (Fig. 5f-g; Extended Data Fig. 5c). Together, these results indicate that FER-mediated CWI monitoring is required for phellem differentiation in response to mechanical stress (Fig. 5i).

## DISCUSSION

The transition from endodermis to phellem is essential to ensure continuous protection from the external environment during secondary growth. After secondary growth initiation, we show here that pericycle cells acquire phellem identity, yet identity alone is insufficient to trigger differentiation. Our data supports a model in which rupture of the overlying endodermis releases mechanical constraints on underlying properiderm cells, leading to their expansion and phellem differentiation (Fig. 5i). Consistent with this model, local reduction of mechanical constraints, either through hyperosmotic treatment or targeted endodermal ablation, was sufficient to induce cell expansion and phellem differentiation in properiderm cells beneath the affected endodermal cells. A similar mechanism has been described in leaves, where removal of outer tissues releases pressure on underlying mesophyll cells, allowing their expansion and activation of the epidermal fate regulator *ATML1* to drive epidermal regeneration^46^.

Our results show that expansion of properiderm cells following endodermis rupture is the first physical response to mechanical release, but expansion alone is insufficient to trigger differentiation. Phellem differentiation requires FER-dependent signalling through an intact and mechanically competent wall (Fig. 5i). In *fer-4* mutants, properiderm cells expand normally but fail to deposit lignin and suberin, indicating that expansion alone is insufficient to drive differentiation and that FER-mediated mechanosignalling is required for phellem differentiation. Pectin-mediated wall elasticity appears particularly important for phellem differentiation, as perturbing pectin composition strongly reduced differentiation. This may reflect impaired FER-mediated mechanosensing, consistent with FER’s ability to bind to de-methylesterified pectins to transduce mechanical signals in other contexts^47^. Further work is needed to confirm this link in the context of phellem differentiation. Since *fer-4* mutants did not completely lack phellem differentiation, additional mechanosensing pathways^48,49^ possibly act in parallel to FER signalling to initiate phellem differentiation.

Our data show that phellem differentiation proceeds through three distinct stages. Mature phellem exhibits uniform suberin deposition and polarized lignin accumulation forming the characteristic tonsure pattern. This lignin pattern resembles the polar lignin cap of the exodermis^50^, but in phellem the outer tangential wall is only partially lignified. Unlike the exodermis, which overlies non-expanding tissues, phellem surrounds tissues that continue to expand during secondary growth. We propose that partial lignification in the tangential wall likely provides mechanical flexibility, allowing barrier cells to accommodate underlying expansion while maintaining barrier integrity.

In mature roots, wounding triggers phellem regeneration through a gas diffusion-mediated mechanism^9^. In this study, endodermis genetic ablation suggests that direct environment contact is not required for phellem differentiation; however, ablation may compromise the apoplastic diffusion barrier function, permitting gas diffusion normally restricted by the endodermis^51^, leaving the contribution of gaseous signals in this context unresolved. Beyond gas-mediated regulation, suberization in both endodermis and mature phellem dynamically responds to nutrient availability, water status, salinity and temperature^28,52,53^. Here, we further identify mechanical cues as an instructive signal for phellem differentiation. Thus, it is likely that multiple context-dependent signals orchestrate phellem differentiation, ensuring the formation of a robust, continuous and adaptable barrier.

## METHODS

### Plant materials and growth conditions

*Arabidopsis thaliana* ecotype Col-0 was used as the wild-type. *pPER15::erVenus*, *pPER49::erVenus*, *shr-2*, *scr-4*, *myb36-2;sgn3-3*, *gelpQ*, *abi1-1*, *pyr1;pyl1,2,4,5,8*, *fer-4*, *herk2*, *the1-1*, *the1-4*, *the1-1;herk2* and *the1-1;fer-4* have been described previously^12,19,22–24,30,31,41–45^.

For all experiments, Arabidopsis seeds were surface-sterilized in 70% ethanol for 5 minutes and rinsed twice with sterile water. Sterilized seeds were stratified at 4°C in darkness for at least 2 days. Seeds were plated on half-strength Murashige and Skoog (MS) media (Duchefa) plates containing 0.05 % MES, 1 % agar and 1 % sucrose (pH5.8). The age of the seedlings was determined from the time they were positioned vertically in the growth chamber set to 22°C under long-day conditions (16h of light and 8h of dark).

### Cloning

*pCASP1::XVE>>m-BAX-3xYFP* was generated from *pCASP1::XVE/*pDONR, *m-BAX*(-)stop/pDONR (Zhang et al. 2025), *3xYFP-OcsT*/pDONR, pFRm43GW by MultiSite Gateway LR clonase reaction. The generated plasmid was introduced into Col-0.

### Surgical injury of the endodermis and chemical treatments

Four-day-old seedlings were used for surgical injury experiments. A longitudinal incision was made within 3 mm below the root-hypocotyl junction using a needle under a dissection microscope by gently pulling cotyledons upward to create tension in roots. Wounded roots were imaged at 3, 6, 24, 48 and 72 hai for confocal microscopy and TEM analysis. Only sections where the wound reached the endodermis without ablating pericycle cells were analysed.

For XVE-based gene induction, seeds were sown on MS-plates containing 5 µM 17-β-oestradiol (Sigma), as a synthetic oestrogen derivative, or an equivalent volume of DMSO as a mock treatment. The 17-β-oestradiol stock solution (20 mM) was prepared in DMSO and stored at −20°C.

For hyperosmotic stress, three-day-old seedlings were transferred to MS-plates containing 400 mM sorbitol (Duchefa). For ABA treatment, a 100 mM stock solution of ABA (Duchefa) was prepared in ethanol and diluted to a 1 µM or 5 µM working concentration. Three-day-old seedlings were transferred to the ABA-containing MS-plates. Mock treatment consisted of growth medium with the same solvent without active compound.

Ten-day-old wild-type seedlings were transferred into MS-plates containing 0.1 µM isoxaben (ISX; Fluka analytical) and 50 µM EGCG (MedChemExpress), prepared from 10 mM stock solution in DMSO to alter cell wall integrity.

### Histological sectioning and staining

Samples were fixed in 4% paraformaldehyde in 1xPBS for 15-30 minutes, followed by two washes in 1xPBS. The samples were embedded in 5% agarose in 1xPBS and sectioned at 200-µm-thick cross-sections using a vibratome.

For staining the cell wall, sections were stained in 1xPBS with 1 µl/ml Renaissance SCRI 2200 (SR2200; Renaissance Chemicals). Lignin staining was performed using ClearSee containing 50 µg/ml Basic Fuchsin (Sigma-Aldrich) for 2 hours, followed by washing with ClearSee until no residual stain remained and a final rinse in 1xPBS. Samples were mounted in 1xPBS with 1 µl/ml SR2200 for imaging.

For simultaneous lignin and suberin visualization, sections were stained overnight in ClearSee supplemented with 50 µg/ml Basic Fuchsin, washed with ClearSee and twice with 100% ethanol, then stained in ethanol with 0.01% Fluorol Yellow 088 for 1h. Before imaging, the sections were washed with 1xPBS and mounted with 1xPBS supplemented with 1 µl/ml SR2200.

### Microscopy and data analysis

The live imaging of fluorescence reporters was conducted on the Leica M165 FC fluorescent stereomicroscope equipped with the Leica Application Suite X package.

Fluorescent markers and stains were imaged with the Leica Stellaris 8 confocal microscope (63x objective). Confocal images were obtained with Leica LAS AF Software. All confocal images with multiple channels were imaged in sequential scan mode. Confocal settings may have varied between experiments but always stayed the same for the experimental sample and the respective control.

Fluorescent signal intensity and cell area quantifications were performed using Fiji (v1.54). *PER15* and *PER49* fluorescent intensities from stereomicroscope images were quantified by selecting a region of interest (ROI) 3-5 mm below the hypocotyl. To measure YFP signal intensities or cell area in cross section images, cell wall images were segmented using Fiji plugin. After manual correction, YFP signal intensity with minimum threshold or cell area in each cell was measured based on the segmented images. For Fig. 2c, YFP-positive pericycle cells were defined as those with signal intensities above the certain threshold. Their proportion was calculated relative to the total pericycle cell number in each root. XPP cells were excluded from *PER15* and *PER49* signal intensities measurements (Fig. 1c, Extended Data Fig. 1b–c), properiderm establishment analyses (Extended Data Fig. 1a), and cell area measurements (Fig. 4b–c; Extended Data Fig. 3a), because periderm-specific measurements were unreliable due to the ambiguous boundary between vascular tissues and periderm in the XPP lineage. Stele length was measured as the distance between diagonally opposite XPP cells across the primary xylem.

All statistical analyses were performed using MS Excel (v2508) and RStudio (v.2026.02.0+392). Dataset normality was assessed using the Shapiro-Wilk test. For comparisons between two groups, a two-tailed t-test was applied if data were normally distributed (*P* ≥ 0.05), and a Wilcoxon rank-sum test if not (*P* < 0.05). For comparisons among three or more groups, one-way ANOVA was followed by Bonferroni-corrected pairwise t-test when both groups followed a normal distribution or Bonferroni-Wilcoxon rank-sum test when the assumption of normality was rejected. Categorical data, such as the frequency of phellem differentiation (Fig. 3a and 4f–g), were analysed using Fisher’s exact test with Bonferroni correction. In Fig. 4f–g, the phenotypes “patchy” and “absent” or “non-lignosuberized” and “collapsed” were combined into one category as “not continuous” and “non-lignosuberized”, respectively, for statistical comparison.

### Sample fixation and lignin staining for transmission electron microscopy

Arabidopsis seedlings were fixed in 2.5% glutaraldehyde solution (EMS, Hatfield, PA) in 0.1 M phosphate buffer pH 7.4 for 1 h at room temperature and post-fixed in a fresh mixture of 1% osmium tetroxide (EMS, Hatfield, PA) with 1.5% potassium ferrocyanide (Sigma, St. Louis, MO) in phosphate buffer for 1 h at room temperature. The samples were then washed twice in distilled water and dehydrated in acetone solution (Sigma, St. Louis, MO) at graded concentrations (30%, 40 min; 50%, 40 min; 70%, 40 min; and 100%, 3 × 1 h). This was followed by infiltration in Spurr resin (EMS, Hatfield, PA) at graded concentrations (Spurr 33% in acetone, 6 h; Spurr 66% in acetone, 8 h; Spurr 100%, 2 × 12 h) and finally polymerized for 48 h at 60 °C. 50-nm-thick sections were cut transversally at 2, 3 or 5 mm below the hypocotyl, using a Leica UC7 (Leica Mikrosysteme), picked up on a copper slot grid 2 × 1 mm (EMS, Hatfield, PA) coated with a polystyrene film (Sigma, St. Louis, MO). Ultrathin sections were post-stained with 1% of permanganate potassium (KMnO4) in H2O (Sigma, St Louis, MO, US) for 45 min and rinsed several times with H2O. Micrographs were taken with a transmission electron microscope Talos 120Kv (Thermo Fisher Scientific Inc., US) at an acceleration voltage of 120kV with a CETA digital camera (Thermo Fisher Scientific Inc., US). Panoramic were acquired using MapsTM software (Thermo Fisher Scientific Inc., US) and aligned with the software IMOD^54^.

## ACKNOWLEDGEMENTS

We thank R. Sozzani, L. Bacete and O. Hamant for providing published seeds; M. Herpola, K. Laine, W. Virtanen and A. Huttunen for technical assistance and the Light Microscopy Unit (University of Helsinki) for the confocal microscopy equipment and technical assistance; and B. Wybouw, M. Lyu, W. Fatz, J. Sirén, J. Vainio, M. López-Sancho and L. Bacete for feedback on the manuscript. This work was supported by grants from the European Research Council (ERC CoG CORKtheCAMBIA 819422 to J.L.-O., H.I. and A.P.M.); the Research Council of Finland (grant number 364181 to J.L.-O., H.I., and A.P.M.); the University of Helsinki (Doctoral Programme in Plant Science to J.L.-O.); the EMBO Postdoctoral Fellowship (ALTF 128-2020 to H.I.); the Japan Society for the Promotion of Science (Overseas Research Fellowships to H.I.), the Swiss National Science Foundation (SNSF, 310030-197739 and 320030-231593 to N.G.).

## AUTHOR CONTRIBUTION

J.L.-O. and A.P.M. conceived the project; J.L.-O., H.I., J.A.-S. and A.P.M. designed the experiments; J.L.-O. performed all the experiments and its analysis, except for the surgical ablation imaging, which was performed by H.I. and quantified by J.L.-O.; N.G. supervised the TEM microscopy imaging, which was performed by D.D.B., J.L.-O., and E.B.; J.L.-O. analysed all the TEM data; J.L.-O. wrote the first draft of the manuscript; J.L.-O., H.I., and A.P.M. wrote the manuscript with input from all the authors.

## COMPETING INTERESTS

The authors declare no competing interests.

## DATA AVAILABILITY

All the data that support the findings of this study are shown in this article and Extended Data Figures. Material requests should be directed to the corresponding authors.

## Notes

### Competing Interest Statement

The authors have declared no competing interest.

