## Supplementary Figures for "Mechanical cues trigger phellem differentiation during barrier transition"

### Extended Data Figures

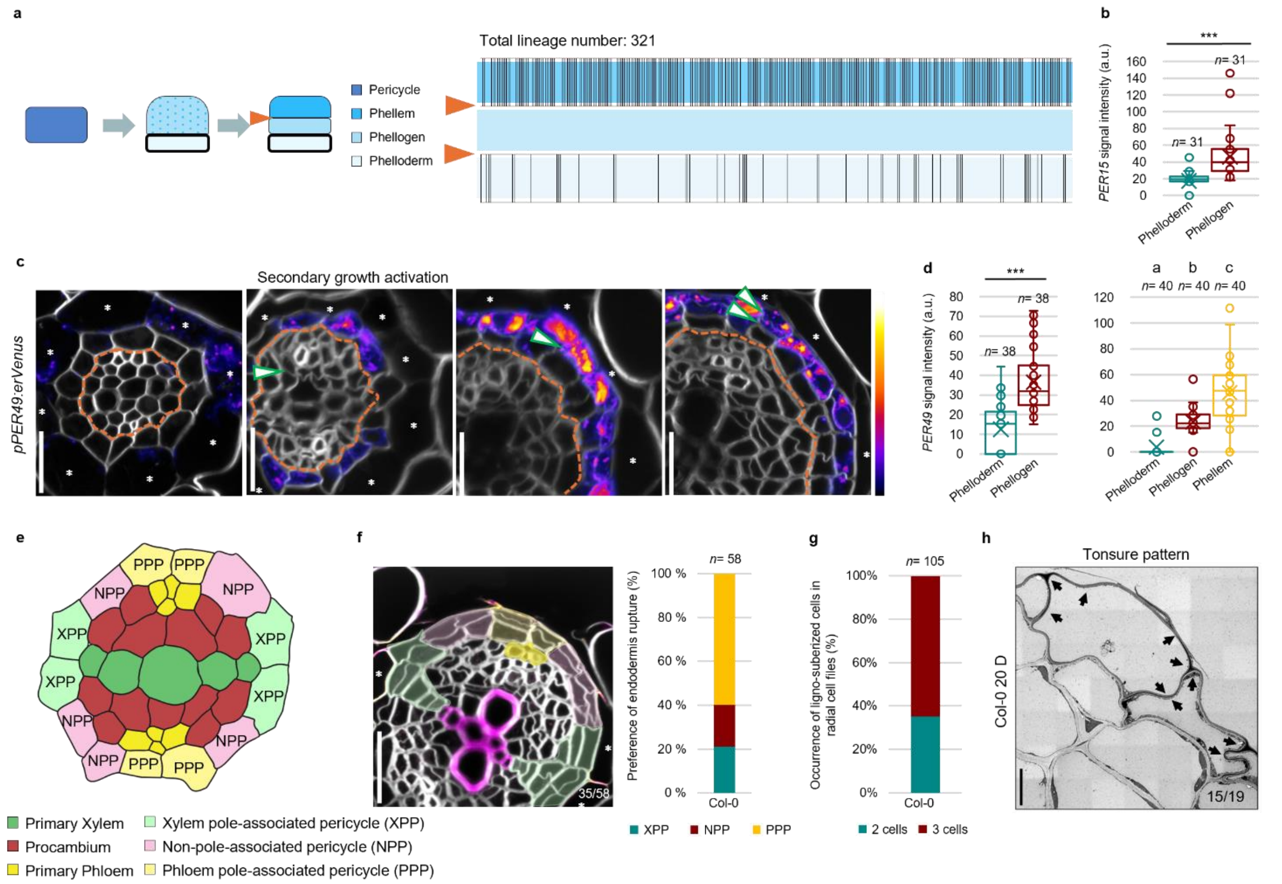


**Extended Data Fig. 1 | Establishment of periderm tissues, phellem identity acquisition and persistence of the lignin tonsure pattern in mature phellem.**

**a**, Schematic of properiderm establishment (left) and quantification of the thinnest cell wall position within a three-cell radial file (right). Newly formed cells (black bars) are plotted according to the position of the thinnest cell wall (orange arrows). **b,** Quantification of *PER15* signal intensity in a two-cell radial file. Two-tailed Wilcoxon rank-sum test (***; $P < 0{,}001$). **c,** Confocal microscopy of the *pPER49::erVenus* reporter line across developmental stages. Venus signal intensities are shown according to the colour map on the right. **d,** Quantification of *PER49* signal intensity in a two- or three-cell radial files (left and right, respectively). Two-tailed Wilcoxon rank-sum test for pairwise comparisons (***; $P < 0{,}001$); one-way ANOVA followed by Bonferroni-corrected two-tailed Wilcoxon rank-sum test were used for three groups (different letters indicate significant differences between groups; $P < 0.001$). In **b** and **d**, boxes represent the median values and interquartile range (IQR), whiskers extend to 1.5×IQR, x marks the mean, and the circles indicate measurements from individual cells. n indicates the number of examined cells in **b** and **d**. **e,** Schematic of pericycle cells classified according to their position relative to the xylem and phloem. **f,** Confocal microscopy of wild-type roots showing a regional preference of endodermis rupture in the PPP lineage position (left) and quantification of the regional preference of endodermis rupture (right). Fractions on the panel indicate the proportion of roots displaying the similar endodermis rupture shown in the image. n indicates the number of examined roots. **g,** Quantification of ligno-suberized cells in radial cell files after native endodermis rupture. n indicates the number of examined lineages. **h,** TEM cross-section of a 20-day-old root showing the lignin tonsure pattern in the mature phellem. Fractions on the panels indicate the proportion of cells exhibiting similar secondary cell wall patterns shown in the images. Endodermal cells are marked with asterisks in **c** and **f**. Cell walls (CW) were stained with SR2200 (grays); lignin with Basic Fuchsin (magenta); and suberin with Fluorol Yellow (yellow). The experiments in **a**, **f** and **g** were independently repeated five times; **b-d** were repeated three times independently, and **h** were obtained from one root. Scale bars: 20 µm (**c**), 10 µm (**f**), 5 µm (**h**).

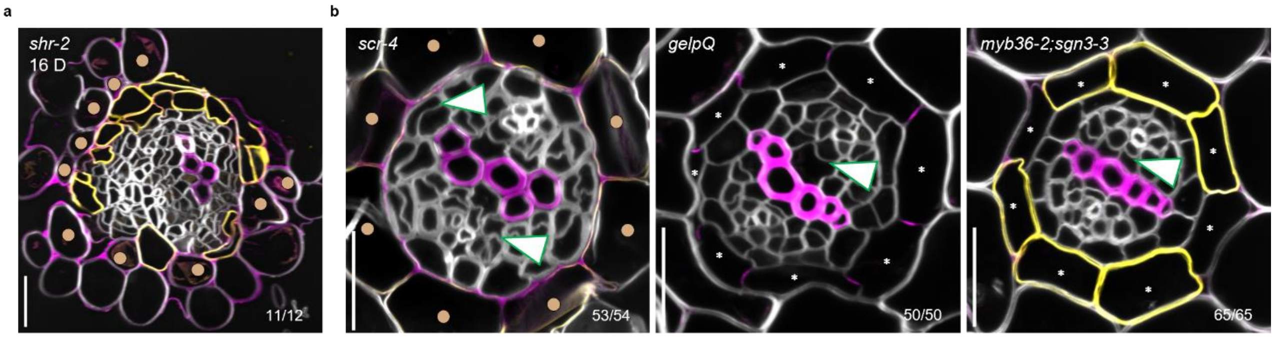


**Extended Data Fig. 2 | Endodermis identity and barrier function are not required to inhibit phellem differentiation.**

**a**, Confocal microscopy of 20-day-old *shr-2* roots stained for lignin and suberin. Fraction indicates the proportion of roots showing phellem differentiation. **b,** Confocal microscopy of endodermis mutant roots with impaired identity (*scr-4*) or barrier function (*gelpQ* and *myb36-2;sgn3-3*) after secondary growth activation, stained for lignin and suberin. Premature pericycle cell divisions were observed in *scr-4* roots. Fraction indicates the proportion of roots lacking secondary cell wall deposition in pericycle cells. Green arrowheads indicate periclinal division. Light brown dots mark the mutant layer in *shr-2* and *scr-4*. Cell walls were stained with SR2200 (white); lignin with Basic Fuchsin (magenta); and suberin with Fluorol Yellow (yellow). Experiments in **a** and **b** were independently repeated three times. Scale bars: 20 µm (**a**,**b**).

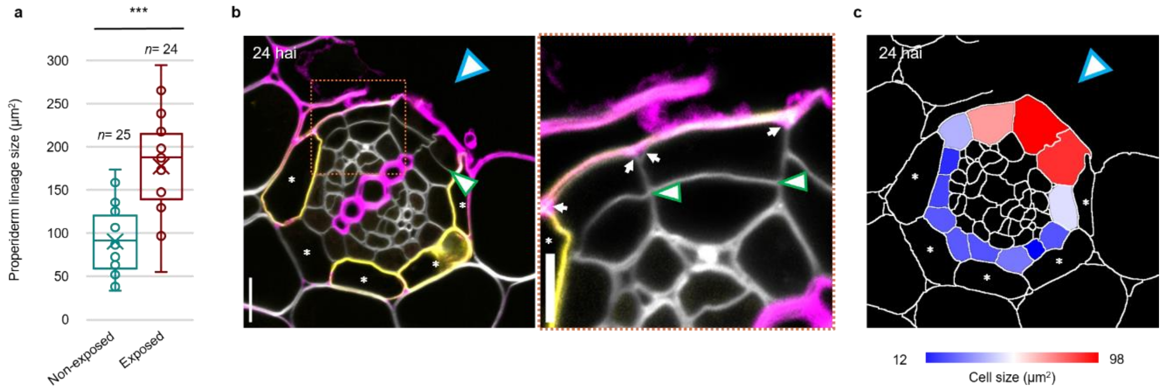


**Extended Data Fig. 3 | Endodermis rupture leads to expansion and phellem differentiation of exposed properiderm.**

**a**, Quantification of non- and exposed properiderm lineage during native endodermis rupture. Boxes represent the median values and interquartile range (IQR), whiskers extend to 1.5×IQR, x marks the mean, and the circles indicate measurements from individual lineages. Two-tailed Welch's t-test was used to compare non-exposed properiderm to exposed containing differentiated phellem (***; $P <$ 0,001). n indicates the number of examined lineages. **b-c**, Surgically cut wild-type roots at 24 hai. **b**, Confocal microscopy showing lignin and suberin depositions and periclinal cell divisions (green arrowhead). Close-up of highlighted region in the right panel showing the lignification pattern (white arrows). **c**, Heatmap of pericycle-derived cell size at 24 hai. Endodermal cells are marked with white asterisks and the cut site in blue arrowheads in **b**,**c**. Cell walls were stained with SR2200 (white); lignin with Basic Fuchsin (magenta); and suberin with Fluorol Yellow (yellow). Experiments in **a-c** were independently repeated three times. Scale bars:10 µm (**b**), 5 µm (**b**, close-up).

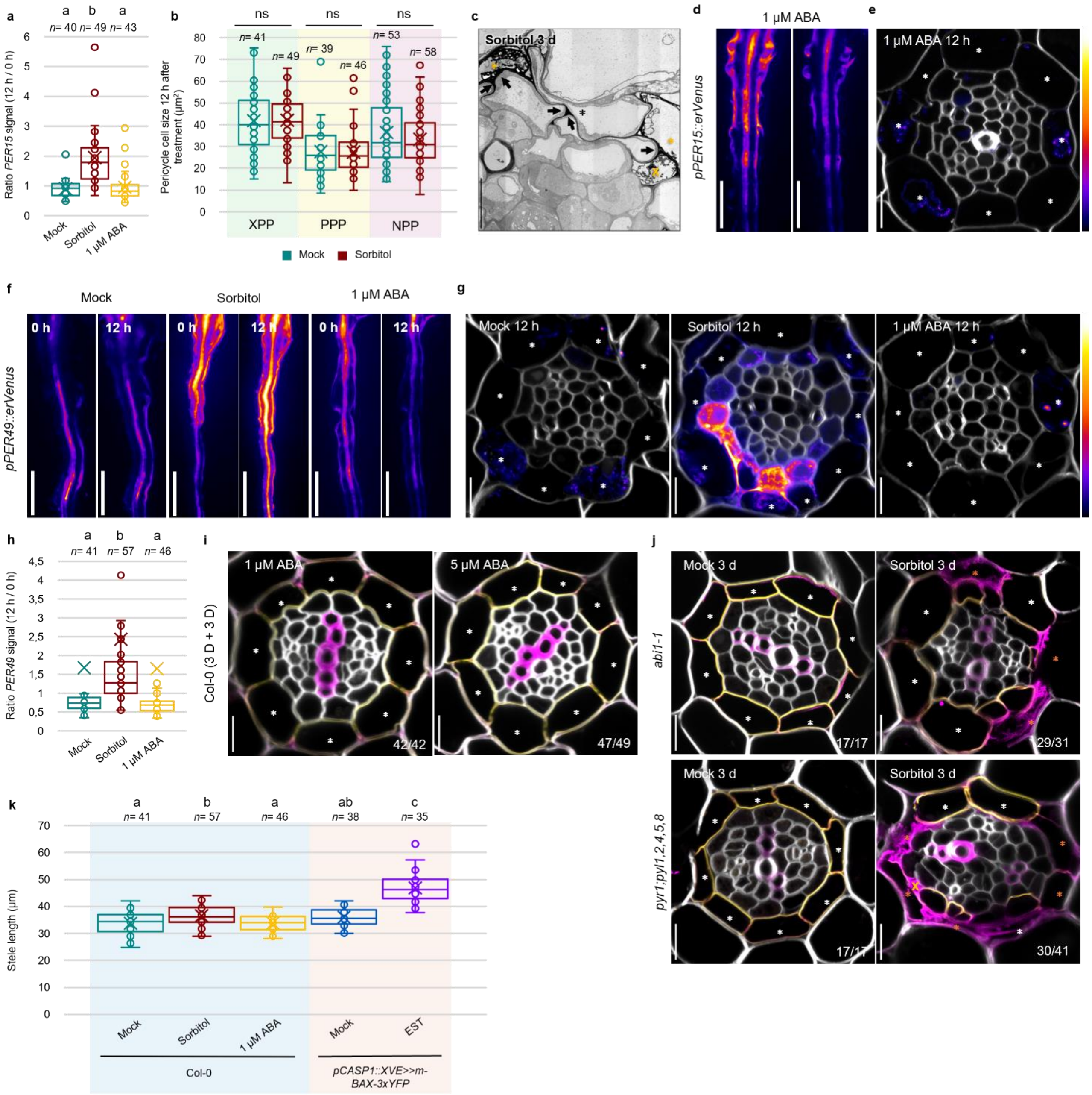


**Extended Data Fig. 4 | Hyperosmotic stress induces endodermal collapse and promotes phellem differentiation independently of ABA signalling.**
**a**, Quantification of *pPER15::erVenus* signal (12 h / 0 h) after mock, sorbitol or ABA. **b**, Pericycle cells area 12 h after mock or sorbitol treatment. **c**, TEM image showing the lignin patterning (black arrows) and cell collapse in endodermis (orange asterisks) and pericycle (orange cross). **d-e**, *pPER15::erVenus* seedlings treated with mock or 1 µM ABA for 12 h, showing *PER15* signal by stereomicroscopy (**d**) and confocal microscopy (**g**). **f-g**, *pPER49::erVenus* seedlings treated with mock, sorbitol or 1 µM ABA for 12 h, showing *PER49* signal under the stereomicroscopy (**f**) and confocal (**g**) . **h**, Quantification of *PER49* signal (12 h / 0 h). **i**, Confocal microscopy of wild type roots 3 days after 1 or 5 µM ABA treatment showing lignin and suberin patterning. **j**, Confocal microscopy of ABA-insensitive mutants after mock or sorbitol treatment for 3 days. Orange crosses indicate collapsed pericycle cells, and collapsed endodermal cells are indicated with orange asterisks.

Fractions in **i-j** denote the frequency of the presented phenotype in independent roots. **k**, Stele length in mock-, sorbitol- or ABA-treated wild-type roots and mock- or EST-treated *pCASP1::XVE>>m-BAX-3xYFP* roots. Boxes represent the median values and interquartile range (IQR), whiskers extend to 1.5×IQR, x marks the mean, and the circles indicate measurements from roots (**a,h,k**), cells (**b**). In **a** and **h**, One-way ANOVA test followed by Bonferroni-corrected Wilcoxon test was used ($P < 0.001$). In **b**, two-tailed Welch's t-test was used to compare XPP cell size**;** and two-tailed Wilcoxon rank-sum test was used to compare PPP and NPP cell size (ns, $P > 0{,}05$). One-way ANOVA followed by Bonferroni-corrected pairwise Welch's t-tests was used in **k**. Intact endodermal cells are marked with white asterisks in **a**,**d**,**g**,**h**,**l** and in black asterisks in **i**. n indicates the number of examined roots (**a**,**h**,**k**) or cells (**b**). Collapsed endodermal cells are indicated with orange asterisks in **h-i**,**l**. Cell walls were stained with SR2200 (white); lignin with Basic Fuchsin (magenta); and suberin with Fluorol Yellow (yellow). Experiments in **a-b** and **d-j** were independently repeated three times; Col-0 samples in **k** were repeated three times, while the experiments using *pCASP1::XVE>>m-BAX-3xYFP* were repeated four times, and **c** were obtained from one root. Scale bars: 1 mm (**d**,**f**), 10 µm (**e,g**,**i-j**), 5 µm (**c**).

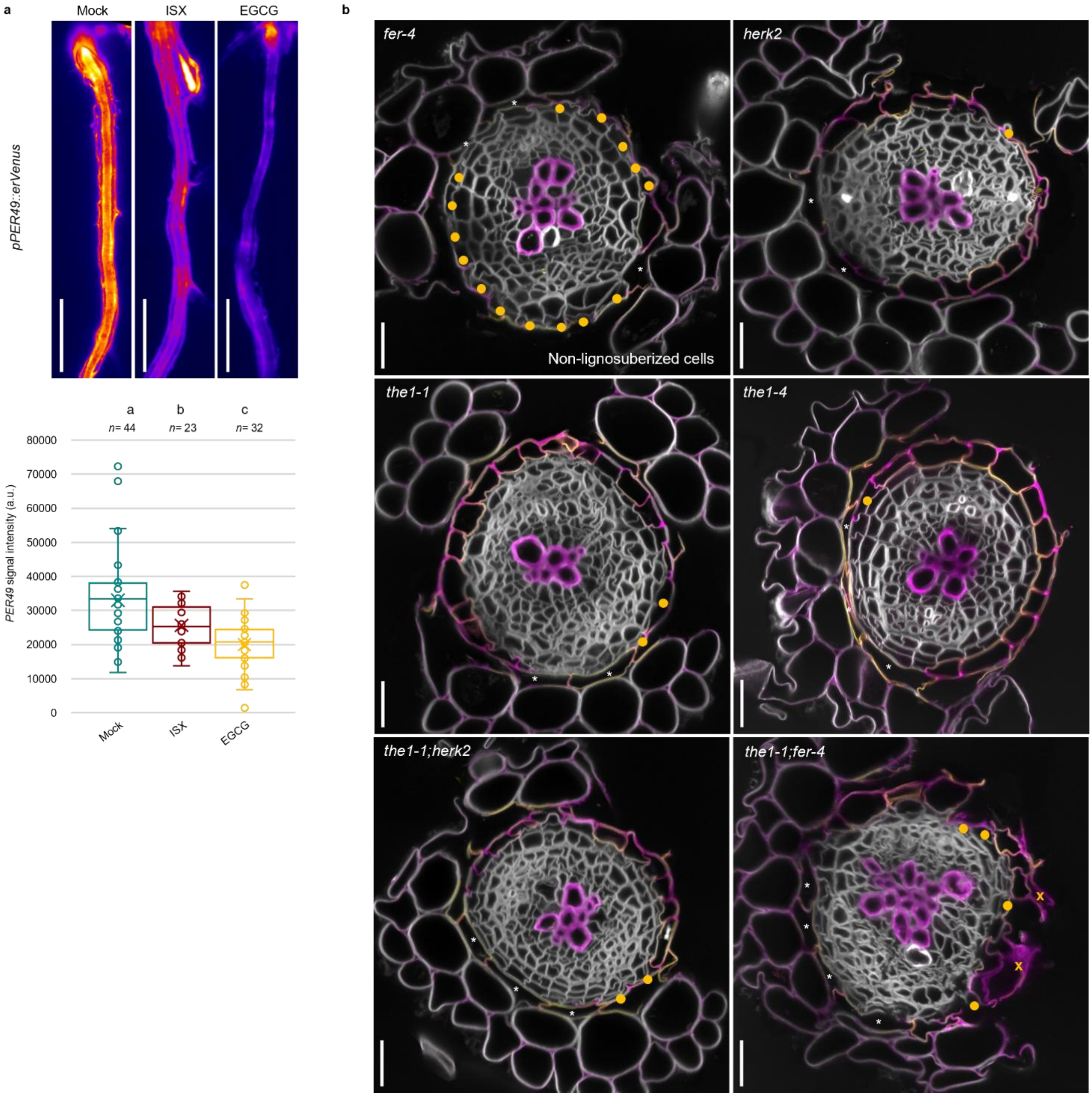


**Extended Data Fig. 5 | CWI receptor mutants reveal a requirement of FER-dependent mechanical signalling during phellem differentiation.**

**a-b**, *pPER49::erVenus* seedlings treated with mock, ISX or EGCG for 24 h during the transition stage, showing *PER49* signal by stereomicroscopy (**a**) and the quantification of the overall *PER49* signal intensity (**b**). Boxes represent the median values and interquartile range (IQR), whiskers extend to 1.5×IQR, x marks the mean, and the circles indicate measurements from individual roots. One-way ANOVA test followed by Bonferroni-corrected Wilcoxon test was used to compare signal in different treatments; different letters indicate significant differences between groups ($P < 0.05$). n indicates examined roots. **c**, Confocal microscopy of 11-day-old CWI mutant roots at the barrier transition stage, stained for lignin and suberin. Images show representative examples of the most common phenotype of exposed pericycle-derived cells. Orange dots mark undifferentiated phellem exposed cells; orange crosses indicate collapsed cells. Endodermal cells are marked in asterisks. Cell walls were stained with SR2200 (white); lignin with Basic Fuchsin (magenta); and suberin with Fluorol Yellow (yellow). Experiments in *fer-2*, *herk2*, *the1-1* and *the1-1;herk2* were independently repeated six times, whereas *the1-4* and *the1-1;fer-4* were repeated four times. Scale bars: 1 mm (**a**), 20 μm (**c**).
